# Protected areas network is not adequate to protect a critically endangered East Africa Chelonian: Modelling distribution of pancake tortoise, Malacochersus tornieri under current and future climates

**DOI:** 10.1101/2020.08.24.264796

**Authors:** Abraham Eustace, Luíz Fernando Esser, Rudolf Mremi, Patrick K. Malonza, Reginald T. Mwaya

## Abstract

While the international pet trade and habitat destruction have been extensively discussed as major threats to the survival of the pancake tortoise (*Malacochersus tornieri*), the impact of climate change on the species remains unknown. In this study, we used species distribution modelling to predict the current and future distribution of pancake tortoises in Zambezian and Somalian biogeographical regions. We used 224 pancake tortoise occurrences obtained from Tanzania, Kenya and Zambia to estimate suitable and stable areas for the pancake tortoise in all countries present in these regions. We also used a protected area network to assess how many of the suitable and stable areas are protected for the conservation of this critically endangered species. Our model predicted the expansion of climatically suitable habitats for pancake tortoises from four countries and a total area of 90,668.75 km^2^ to ten countries in the future and an area of 343,459.60 - 401,179.70 km^2^. The model also showed that a more significant area of climatically suitable habitat for the species lies outside of the wildlife protected areas. Based on our results, we can predict that pancake tortoises may not suffer from habitat constriction. However, the species will continue to be at risk from the international pet trade, as most of the identified suitable habitats remain outside of protected areas. We suggest that efforts to conserve the pancake tortoise should not only focus on protected areas but also areas that are unprotected, as these comprise a large proportion of the suitable and stable habitats available following predicted future climate change.

## INTRODUCTION

Over the past few decades, there has been growing interest in species distribution models (SDMs) as fundamental tools for the studies of ecology, biogeography, and biodiversity conservation [1]. These models are used to enhance understanding of the factors that alter species distribution, which is critical for adjusting or designing appropriate conservation strategies and for predicting geographical distribution under current and future climatic scenarios [1–3]. Such adjustments and predictions are necessary because climate change poses a severe threat to the conservation of natural landscapes and species across the globe and is reported to be among the primary drivers of the current loss of global biodiversity [4–6]. Climate change has also been reported to accelerate shifts in range extension and the shrinkage of some species [3].

Tropical environments are widely recognized as biodiversity regions with ideal climatic conditions for the survival of a large proportion of reptile species. However, reptiles are currently facing severe threats because of climatic changes [3,7]. Climate change affects reptile biodiversity directly by altering reptile distribution patterns [3,4] and indirectly by threatening conservation areas, making them less habitable for reptile species [8]. For instance, Meng et al. [7] have reported that out of the 274 Tanzania reptile species they studied, 71% (194 reptile species) are vulnerable to climate change, suggesting a significant impact from climate on reptilian diversity. In a different study, predictions about the environmental responses of reptiles to future climatic conditions made using SDMs showed that four endemic Moroccan reptilian species are highly vulnerable to extinction in Morocco if climatic disturbance prevails as predicted [3]. The same study concluded that reductions in species-rich areas were also likely in future climatic scenarios [3].

Like other reptiles, *Malacochersus tornieri* hereafter referred to as the pancake tortoise, is not immune to the effects of climate change. The pancake tortoise is a small, soft-shelled, dorsoventrally flattened chelonian with discontinuous distribution in the scattered rocky hills and kopjes of the savannas of south-eastern and northern Kenya and northern, eastern, and central Tanzania [9–12]. The presence of pancake tortoises has also been reported in northern Zambia [13]. The areas in which pancake tortoises can be found are typically semi-arid; these areas are classified as having a dry climate, corresponding to both Zambezian and Somalian biogeographic regions, according to Linder et al. [14]. The Zambezian biogeographical region is a wider biogeographical region, spreading across Africa from Namibia to Tanzania, while the Somalian biogeographic region is considered a refugium for arid-adapted plants and a centre of endemism for reptiles [15,16].

Although the international animal trade and habitat destruction have been cited as the major threats to the survival of the pancake tortoise [10,11,17,18], the impact of climate change on the species remains largely unknown. Although the IUCN has identified climate change as one of the threats to pancake tortoise populations [10,11], to our understanding, there is no study that has assessed the impact of climate change on the future distribution pattern of pancake tortoises. The IUCN’s *Guidelines for Re-Introductions and Other Conservation Translocations* [19] has pointed out the need to understand and match the current and/or future climate of the destination area as a key climate requirement for introduced/translocated species. Considering that the pancake tortoise is critically endangered [9–12] and listed in the *CITES Appendix II* [13], understanding current and future climatic habitats suitable for this species could be an essential step in charting out a species conservation plan. Therefore, in this study, we used species distribution modeling (SDM) to determine current and future climatic habitats suitable for the pancake tortoise. We chose this method because SDM is the most widely accepted method of predicting climatically suitable habitats [20], which helps to avoid uncertainties in selecting areas for translocation while providing a higher chance of success [19–21].

While protected areas remain an essential approach for conserving biodiversity and protecting it against human-mediated threats [7,22], there are endangered species that inhabit areas outside of protected lands [23]. For example, 14.00% of threatened mammal species, 19.80% of birds, 10.10% of turtles and 26.60% of amphibians inhabit areas outside of the global protected area network [23]. Similarly, the majority of the pancake tortoise population in Kenya occupies habitats outside of protected areas [10]. With time, the ongoing impacts of climate change are expected to inflict changes to suitable habitats for pancake tortoises both within and outside of protected areas [8,24,25]. Therefore, understanding whether these protected areas will continue to be viable for protecting suitable habitats for pancake tortoises in the event of climate change is crucial to the development of specific and appropriate management and conservation plans for the species. Considering the species’ range varies under different climatic scenarios [1,2,26,27] while the size of most protected areas tends to remain the same [28], more species may eventually be placed at risk of extinction, especially threatened species [29]. Therefore, in order to align protected areas with suitable habitat ranges [30] and enhance the conservation of threatened species in different climatic scenarios [28], SDMs are essential.

SDMs have been used to assess the impact of climate change on the distribution of different species (e.g. [24,25,28,31]). These models use location data and environmental variables to predict the suitable distributional range of a species under climate change conditions [24,28], which is essential when designing adequate species management programmes, as well as for endangered species conservation planning [32]. Although Bombi et al. [33] have used SDMs to model the distribution of all African tortoise species, including the pancake tortoise, their study did not predict the future distribution of the species. In this study, we used SDM to assess the distribution of pancake tortoises under current and future climatic conditions and to investigate how much of the climatically suitable habitat occurs within the Protected Areas Network in the Somali-Maasai and Zambezian biogeographical regions. Specifically, we assessed (i) the current and future climatically suitable areas for pancake tortoises, (ii) the occurrence of more stable areas over time and (iii) whether the protected areas will be viable for the conservation of the species. The findings of this study could play a significant role in promoting the use of species distribution studies on species management approaches [34], including the proposal of suitable areas for translocation [20,35,36] and the establishment of nature reserves where species can be protected with minimal human intervention, as recommended by Malonza [10].

## METHODOLOGY

### Study Area

We predicted current and future climatically suitable habitats for pancake tortoises within the two major biogeographical regions of Africa in which the animal occurs naturally (Fig 1B). These regions are the Somali-Masai Regional Centre of Endemism (Somali-Masai RCE) and the Zambezian Regional Centre of Endemism (Zambezian RCE), both of which fall within the semi-arid climatic belt of eastern-south Africa [37,38]. The Somali-Masai RCE covers approximately 1.87 million km^2^ of arid savannah, extending from north-eastern Somalia to the north-eastern province of Kenya and reaching south through Tanzania into the valley of the Great Ruaha; it ends north of Lake Malawi [14,38,39]. The Somali-Masai RCE harbours approximately 4,500 plant species, of which 31.00% are endemic in the region [37,38]. The dominant vegetation in this region is *Acacia* spp. The Zambezian RCE (3.77 million km^2^) extends in the northeast from the Somali-Masai RCE, and its distribution coincides with the Guinea savannas and woodlands and the Karoo-Namib RCE in the southwest [37,38]. It covers the whole of south-central Africa, from the Atlantic seaboard of Angola to the entirety of Mozambique, Tanzania and the uplands of Kenya and Ethiopia [14,39]. In terms of plant richness, it is more diverse than the Somali-Masai RCE, hosting about 8,500 plant species, out of which 54.00% are endemic in the region [37,38].

**Fig 1:**
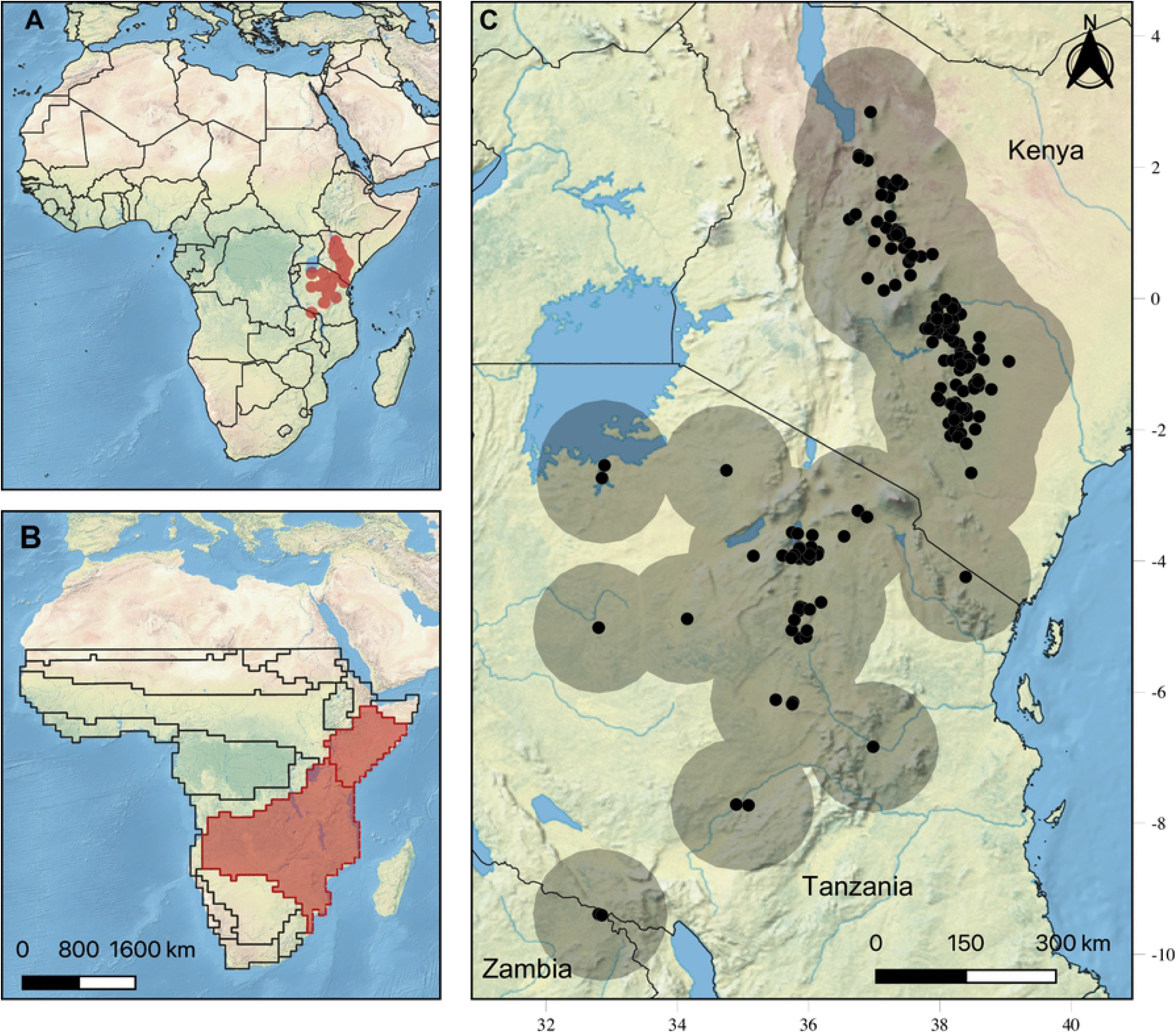
Current natural occurrence of pancake tortoise (*Malacochersus tornieri*) in the sampled areas. (A) Africa showing the buffered area (red) where location data was obtained. (B) Africa showing Zambezian and Somalia biogeographical region (red). (C) Pancake tortoise occurrences (black) with 1-degree wide buffer for each presence record (grey).

### Study Species

The critically endangered pancake tortoise, *Malacochersus* [12], is a monotypic genus endemic to East Africa [10,40]. In East Africa, the pancake tortoise is restricted to Somali-Maasai and Zambezian vegetations [12,18,41–43]. In Tanzania, the species distribution is discontinuously scattered from the south-eastern shores of Lake Victoria to the Maasai Steppe and southward up to Ruaha National Park [10,12,18]. In Kenya, pancake tortoises are disjointly distributed from the northern to southern areas, passing though the central to south-eastern regions of the country [10,12]. In Zambia, the species has been recorded only in the northern Nakonde District that borders Tanzania [12,13]. The preferred micro-habitats for pancake tortoises are kopjes, rock outcrops and rocky hillsides [9,10,18] with an annual rainfall of 250 - 500 mm [42] and an elevational range of 400 - 1,600 m above sea level [44]. From the past two generations to the next generation, the observed and expected population of pancake tortoises is expected to decline by 80.00%, with overexploitation and habitat destruction being the primary drivers [12]. Currently, the IUCN identifies biological resource use (intentional use) and agriculture and aquaculture (small-holder farming), as well as climate change (severe drought), as among the major threats to the habitat and population of pancake tortoises [12].

### Pancake Tortoise Occurrence Data

We obtained occurrence data from the field, online databases and previous studies. On the ground, we collected pancake tortoise location data from eight sites in Tarangire National Park, three sites in the Babati district of the Manyara region, five sites in the Kondoa districts and two sites in Chemba district, both from the Dodoma region in central Tanzania.

We also downloaded pancake tortoise locations from the GBIF (https://www.gbif.org/) and VertNet (http://vertnet.org/). We used the rgbif [45] and rvertnet [46] R packages to download the occurrence data from the GBIF and VertNet, respectively. Both databases were accessed on 5 January 2020, and we downloaded all *Malacochersus tornieri* locations identified in Tanzania and Kenya. We did not find any pancake tortoise occurrences in Zambia in the two databases. From the online databases, we excluded data with absent or incomplete coordinates and duplicate locations as well as non-natural locations, such as tortoise collection points, captive breeding sites and pet-animal release sites. Additionally, we searched for pancake tortoise locality records in the EMYSystem Global Turtle Database [47] and then used Elevation Map (https://elevationmap.net/) to obtain location coordinates.

From previous studies, we extracted the names of the places where pancake tortoises were recorded/observed. For Tanzania, we used sites mentioned by Klemens and Moll [18] as well as point locations collected by Zacarias [48], while for Zambia we used point locations mentioned by Chansa and Wagner [13]. In Kenya, we obtained pancake tortoise sites from Malonza [10] and Kyalo [44]. After obtaining the site names, we used Google Maps (https://www.google.co.tz/maps/), Elevation Map (https://elevationmap.net/) and Mindat (https://www.mindat.org/) to obtain coordinates for each site. If the site was not available online, we contacted individuals currently or previously working in the area in order to obtain coordinates. From all sources, we obtained data for a total of 224 occurrences, with most occurrence points falling within the current IUCN pancake tortoise distribution range (Fig 1C).

### Bioclimatic Variables

Bioclimatic variables were obtained from the CHELSA database [49] with 30 arc-seconds resolution. The modelling domain comprised Zambezian and Somalian biogeographical regions [14]. These regions were selected because they represent the areas where pancake tortoises exist naturally [12,13,18,41–43]. We obtained variables for the two intermediate Representative Concentration Pathways (RCPs), RCP-4.5 and RCP-6.0, for the years 2050 (mean climate between 2041 and 2060) and 2070 (mean climate between 2061 and 2080). These mid-impact RCPs are the most desirable for future conservation planning, since they present a more realistic path compared to the extreme RCPs (2.6 and 8.5) which may incorporate too many uncertainties, causing projections to be unreliable. Variations in future scenarios were also accessed through ten Global Circulation Models (GCMs) available in CHELSA; we avoided those with high co-dependency [50], resulting in the selection of MIROC5, CESM1-CAM5, IPSL-CM5A-MR, FIO-ESM, GISS-E2-H, CSIRO-Mk3-6-0, GISS-E2-R, GFDL-ESM2G, MIROC-ESM-CHEM and MRI-CGCM3.

Variables were first submitted to a visual analysis, in which we deleted both the precipitation of the warmest quarter (BIO 18) variable and the precipitation of the coldest quarter (BIO 19) variable due to statistical artifacts, that may not represent the continuous gradient of reality, in the study region. Those artifacts are generated due to a difference in which quarter is the warmest (e.g. BIO18), causing the precipitation of one cell to be the sum from January-February-March, while the very next cell is the summed precipitation from February-March-April. We then masked variables with one degree-wide buffer from each presence record (Fig 1A and C) and excluded variables with a high variance inflation factor (VIF > 3) and highly correlated variables (r > 0.7). This left us with six variables: the mean diurnal range (BIO 2), the isothermality (BIO 3), the mean temperature of the wettest quarter (BIO 8), the precipitation of the wettest month (BIO 13), the precipitation of the driest month (BIO 14) and the precipitation seasonality (BIO 15). These six variables were used to calculate the climatic niche of the species. The selection routine was performed using the usdm package [51] in R 3.6.2 [52]. Models were generated with variables at 30 arc-seconds resolution, while the rasters used to project models were upscaled at a factor of 10, resulting in rasters with a resolution of 2.5 arc-minutes.

### Species Distribution Modelling

For the SDMs, we applied an ensemble method using the sdm package [53] in R 3.6.2 [52]. We implemented five algorithms using different approaches, with proper pseudo-absence selection, following Barbet-Massin et al. [54], as follows: MaxEnt, a machine-learning approach, with 1,000 randomly selected pseudo-absences; Multivariate Adaptive Regression Splines, a regression-based approach, with 100 randomly selected pseudo-absences; Multiple Discriminant Analysis, a classification approach, with 100 pseudo-absences randomly selected outside a surface-range envelope; Random Forest, a bagging approach, with 224 pseudo-absences randomly selected outside a surface-range envelope; and BIOCLIM, an envelope approach, with 100 randomly selected pseudo-absences. Algorithms were implemented using standard procedures within the sdm package [53]. Model evaluation was performed with ten runs of a four-fold cross-validation technique (75.00% training and 25.00% test). In each run, we calculated true skill statistics (TSSs) and the area under the receiver operating characteristic (AUC). To build ensemble models for each scenario, and after some pre-analysis, we selected models with TSSs and AUCs higher than the mean plus half the standard deviation. The mean AUC value was 0.958, with a standard deviation of 0.059 and a threshold equal to 0.988. The mean TSS value was 0.861, with a standard deviation of 0.112 and a threshold equal to 0.917. The selected models were binarized using the AUC threshold. This approach avoided the use of subjective thresholds. Ensembles were built as a committee average of binarized rasters. Afterwards, we normalized the resulting rasters. This returned an ensemble in which 1 represents sites where all models agree with presences, 0 represents sites where all models agree with absences and the values in between are subject to uncertainty, where 0.5 represents cells with the highest uncertainty (i.e. half of the models agree with an absence, while the other half agree with a presence). We also built three potential refugees for the species by summing the normalized rasters from the five scenarios (current, RCP-4.5/2050, RCP-4.5/2070, RCP-6.0/2050 and RCP-6.0/2070). Then, we applied three thresholds, which were calculated by extracting all values greater than zero from the raster and obtaining the 90^th^, 95^th^ and 99^th^ quantiles (2.179, 2.850 and 3.930, respectively).

Furthermore, we calculated climatically suitable areas using a weighted method, multiplying the cell’s committee average by the cell area and summing all values within the rasters. This conservative method was intended to consider the uncertainty underlying each cell, as well as the different occupation proportions. We applied this method to all scenarios, as well as, to every country present in the Zambezian and Somalian biogeographical regions [14]. We also masked results from area calculations with the World Database on Protected Areas v. 3.1 polygons [36] to estimate the climatically suitable areas under protection in the regions, countries and scenarios. Area calculations were performed in R 3.6.2 [52].

## RESULTS

Our model demonstrated high performance, with an average AUC of 0.958 (SD = 0.059) and an average TSS of 0.861 (SD = 0.112). Currently, in the Zambezian and Somalian biogeographical regions, the pancake tortoise has a more extensive range in Tanzania and Kenya than in other countries present in the region (Fig 2). Although there is currently no evidence of records of pancake tortoises in Angola and Ethiopia, surprisingly, the model predicted patches of climatically suitable habitats in those countries under the current climatic scenario (Fig 2). Additionally, the model revealed that the current suitable distribution range of pancake tortoises is 90,668.75 km^2^, with Kenya contributing 61.10% of the current total range, followed by Tanzania (30.32%), Ethiopia (5.03%) and Angola (3.55%) (Table 1). Considering future climatic scenarios, the SDM predicted that the pancake tortoise’s suitable habitat would not decrease. This was observed through the expansion of suitable habitats as predicted by RCP 4.5 and RCP 6.0 (Fig 2; Table 1). The model predicted that the current distribution range would expand by 303.95% in the year 2050 and 342.47% in the year 2070 for RCP-4.5 and by 278.81% in the year 2050 and 311.99% in the year 2070 under RCP-6.0 (Table 1). Similar to in the current scenario, Kenya and Tanzania will continue to have a larger suitable area (Fig 2; Table 1) than other countries. However, the distributional range will expand from the current four countries to ten countries in the future (Table 1).

**Fig 2.**
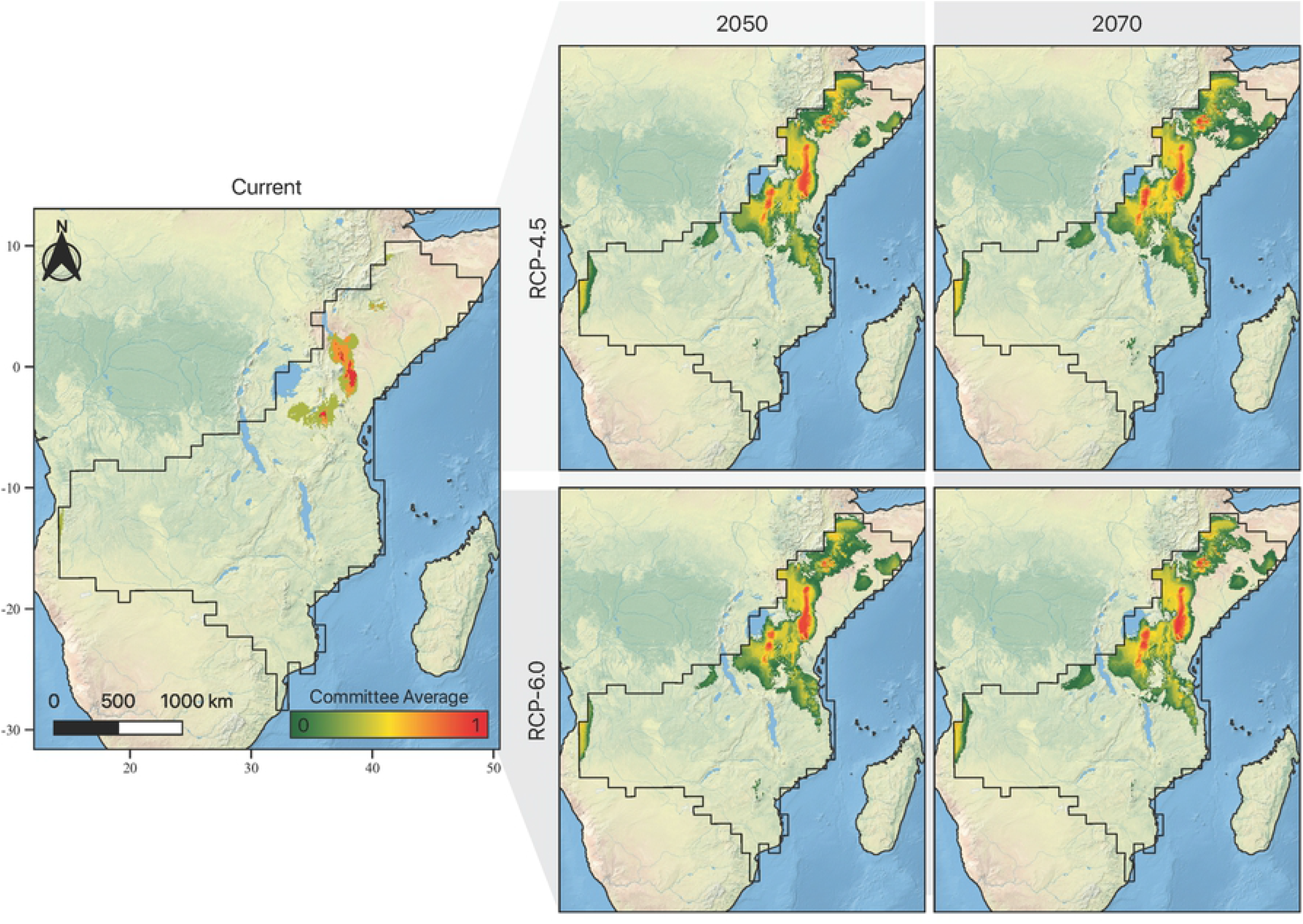
Distribution of pancake tortoise (*Malacochersus tornieri*) in the Somalia and Zambezian biogeographical regions. Current and future (2050 and 2070) climatic suitable habitat for pancake tortoise in the Zambezian and Somalia biogeographical regions considering six bioclimatic variables and two future climate scenarios. Warmer colours show more suitable areas, ranging from red to green.

**Table 1.** *Malacochersus tornieri* suitable area (km^2^) in the countries present in the Zambezian and Somalia biogeographical regions for the current and future climate scenarios.

| Country | Current |  | Future |  |  |  |  |  |  |  |
| --- | --- | --- | --- | --- | --- | --- | --- | --- | --- | --- |
|  |  |  | RCP 4.5 |  |  |  | RCP 6.0 |  |  |  |
|  | Total | Protected | 2050 |  | 2070 |  | 2050 |  | 2070 |  |
|  |  |  | Total | Protected | Total | Protected | Total | Protected | Total | Protected |
| Kenya | 55,401.76 | 19,768.41 | 122,938.04 | 27,285.45 | 129,259.60 | 28,207.22 | 121,659.50 | 27,445.02 | 124,362.9 | 27,954.58 |
| Tanzania | 27,489.20 | 9,268.99 | 136,652.13 | 66,891.79 | 152,800.60 | 73,389.19 | 126,815 | 58,915.30 | 149,008.6 | 71,203.84 |
| Zambia | 0 | 0 | 0 | 0 | 0 | 0 | 0 | 0 | 0 | 0 |
| Mozambique | 0 | 0 | 1,906.56 | 243.10 | 2,424.43 | 397.25 | 996.42 | 64.01 | 1,906.07 | 170.59 |
| Malawi | 0 | 0 | 25.65 | 0 | 34.47 | 0 | 0 | 0 | 38.30 | 0 |
| Zimbabwe | 0 | 0 | 95.31 | 31.78 | 450.11 | 88.55 | 282.20 | 102.28 | 44.29 | 22.14 |
| Angola | 3,219.98 | 0 | 18,528.14 | 222.42 | 19,549.08 | 142.50 | 18,689.62 | 279.76 | 20,321.14 | 333.72 |
| Namibia | 0 | 0 | 0 | 0 | 0 | 0 | 0 | 0 | 0 | 0 |
| Botswana | 0 | 0 | 0 | 0 | 0 | 0 | 0 | 0 | 0 | 0 |
| Somalia | 0 | 0 | 10,233.67 | 0 | 12,181.94 | 0 | 11,891.61 | 0 | 7,692.55 | 0 |
| Ethiopia | 4,557.81 | 311.61 | 71,044.64 | 20,265.41 | 76,881.24 | 20,672.55 | 60,338.92 | 17,527.87 | 65,144.88 | 17,926.38 |
| Democratic | 0 | 0 | 4,283.13 | 29.45 | 6,532.77 | 215.45 | 2,327.59 | 103.07 | 4,388.17 | 61.58 |
| **Republic of** |  |  |  |  |  |  |  |  |  |  |
| **Congo** |  |  |  |  |  |  |  |  |  |  |
| **Burundi** | 0 | 0 | 0 | 0 | 0 | 0 | 0 | 0 | 0 | 0 |
| **Rwanda** | 0 | 0 | 0 | 0 | 0 | 0 | 0 | 0 | 0 | 0 |
| **Uganda** | 0 | 0 | 11.15 | 0 | 31.08 | 0 | 44.59 | 0 | 38.85 | 0 |
| **South Africa** | 0 | 0 | 0 | 0 | 0 | 0 | 0 | 0 | 0 | 0 |
| **TOTAL** | **90,668.75** | **29,349.01** | **366,257.22** | **114,969.40** | **401,179.70** | **123,112.71** | **343,459.60** | **104,437.30** | **373,547.30** | **117,672.83** |
| **% of change** |  |  | **303.95%** | **291.73%** | **342.47%** | **319.48%** | **278.81%** | **255.85%** | **311.99%** | **300.94%** |
| **relative to the** |  |  |  |  |  |  |  |  |  |  |
| **current scenario** |  |  |  |  |  |  |  |  |  |  |

The highly suitable areas (indicated by higher committee averages) are currently present in Kenya and Tanzania (Fig 2), where the species occurs naturally. In the future, highly suitable habitats will expand into Ethiopia as well (Fig 2); however, the species has not yet been recorded in that country. Although there were observations of pancake tortoises in Zambia (Fig 1), our model predicted that the area is not climatically suitable for pancake tortoises in the current and future scenarios (Fig 2). Considering protected lands, we found that a larger suitable habitat for pancake tortoises lies outside of the current Protected Areas Network in both current and future climatic scenarios (Table 1). Currently, 67.63% of the suitable pancake tortoise habitat lies outside of protected areas (Table 1). In the future, the protected suitable area for pancake tortoises will expand from 114,969.40 km^2^ (in 2050) to 123,112.71 km^2^ (2070) in RCP-4.5 and 104,437.30 km^2^ (in 2050) to 117,672.83 km^2^ (in 2070) in RCP-6.0 (Table 1), given the current Protected Area Network. However, the larger suitable habitat of pancake tortoises will continue to lie outside of protected areas in the future (RCP 4.5: 68.61% in 2050 and 69.31% in 2070; RCP 6.0: 69.59% in 2050 and 68.50% in 2070; Table 1).

We identified Kenya, Tanzania, Ethiopia and Angola as the countries that maintain the most stable habitat for pancake tortoises over time (Table 2). However, the highest stability occurs within Kenya, Tanzania and Ethiopia (Fig 3), with only Kenya having a highly stable habitat inside the protected areas (Table 2). The stable habitats for pancake tortoises within the current Protected Areas Network will continue to be smaller than those of habitats in unprotected areas (percentage of stable habitat present in protected areas: less stable [33.08%], average stability [27.97%] and highly stable [14.87%, present in Kenya only]; Table 2).

**Table 2.** Potential climatic stable areas/habitats (in km^2^) for the pancake tortoise per each country present in the Zambezian and Somalia biogeographical regions.

| Country | Less stable |  | Mid stable |  | Highly stable |  |
| --- | --- | --- | --- | --- | --- | --- |
|  | Total | Protected | Total | Protected | Total | Protected |
| **Kenya** | 92,198.02 | 32,556.34 | 57,252.66 | 17,688.00 | 16,066.27 | 2,477.73 |
| **Tanzania** | 50,153.64 | 17,825.94 | 16,299.88 | 4,269.57 | 255.98 | 0 |
| **Zambia** | 0 | 0 | 0 | 0 | 0 | 0 |
| **Mozambique** | 0 | 0 | 0 | 0 | 0 | 0 |
| **Malawi** | 0 | 0 | 0 | 0 | 0 | 0 |
| **Zimbabwe** | 0 | 0 | 0 | 0 | 0 | 0 |
| **Angola** | 2,174.18 | 0 | 83.67 | 0 | 0 | 0 |
| **Namibia** | 0 | 0 | 0 | 0 | 0 | 0 |
| **Botswana** | 0 | 0 | 0 | 0 | 0 | 0 |
| **Somalia** | 0 | 0 | 0 | 0 | 0 | 0 |
| **Ethiopia** | 19,904.86 | 4,004.25 | 8,513.63 | 1,022.46 | 340.53 | 0 |
| **Democratic Republic of Congo** | 0 | 0 | 0 | 0 | 0 | 0 |
| **Burundi** | 0 | 0 | 0 | 0 | 0 | 0 |
| **Rwanda** | 0 | 0 | 0 | 0 | 0 | 0 |
| Uganda | 0 | 0 | 0 | 0 | 0 | 0 |
| South Africa | 0 | 0 | 0 | 0 | 0 | 0 |
| **TOTAL** | **164,430.70** | **54,386.53** | **82,149.83** | **22,980.03** | **16,662.78** | **2,477.73** |

**Fig 3.**
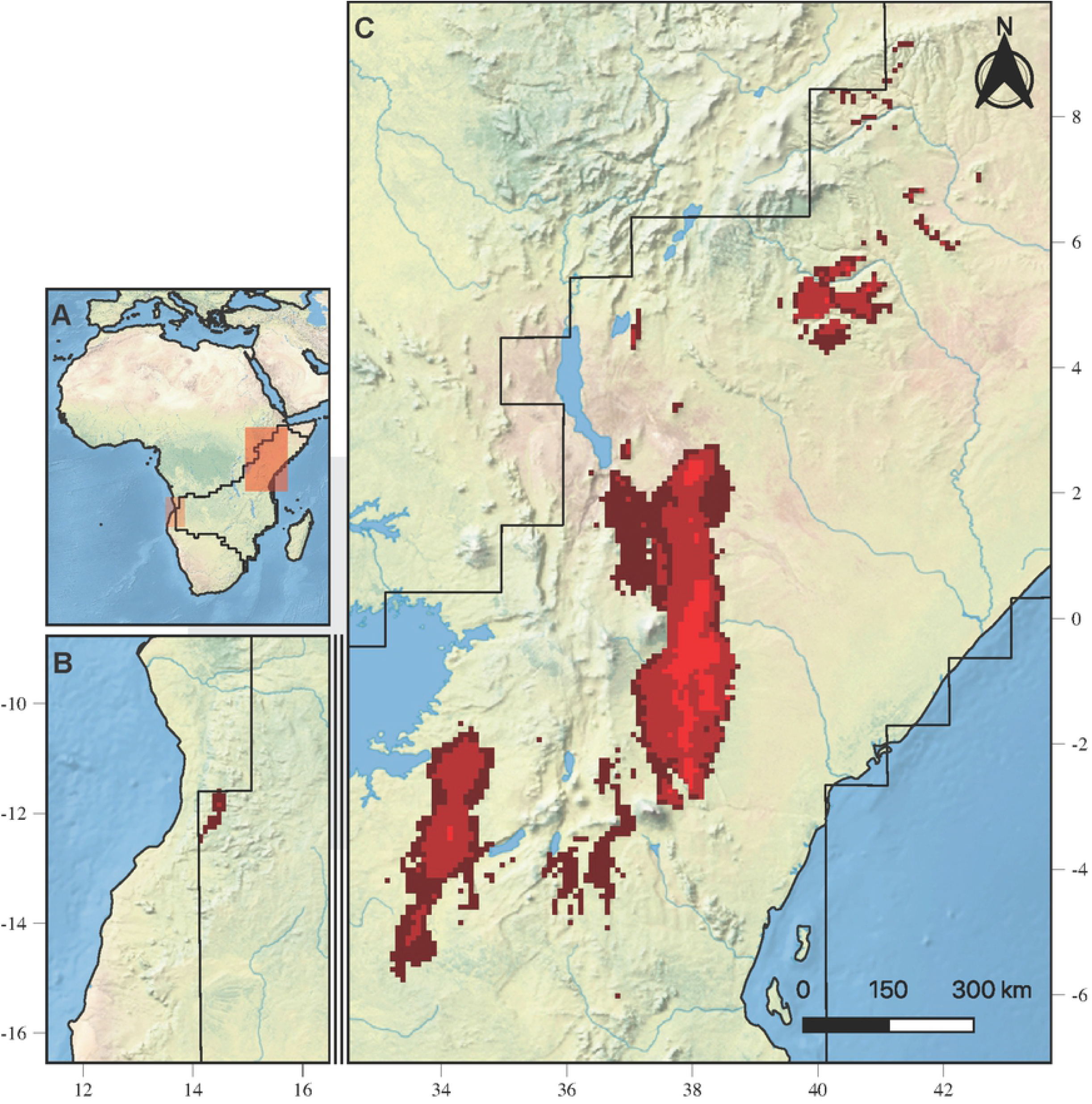
Potential climatic stable areas for the pancake tortoise in the Zambezian and Somalia biogeographical regions. (A) Location of stable areas in Africa (red sheds). (B) Stable areas in Angola. (C) Stable areas in Tanzania, Kenya and Ethiopia. The stable areas were obtained by considering three thresholds from the sum of the five normalized climatic scenarios (current, RCP4.5/2050, RCP4.5/2070, RCP6.0/2050 and RCP6.0/2070). The brighter red colour indicates the more stable site through time.

## DISCUSSION

Our SDM predicted that the suitable climatic habitat for pancake tortoises would be less discontinuously scattered in the Zambezian and Somalian biogeographical regions in the future than in current climatic scenarios (Fig 2). The disjointed distribution of pancake tortoises was also observed in the countries in which they currently exist naturally, which are Tanzania [9,12] and Kenya [10,12]. We further observed that the distributional range of pancake tortoises would expand in the future (Fig 2; Table 1). The expansion of the future distributional ranges of reptiles has also been recorded by Houniet et al. [55] for *Bradypodion occidentale*, González-Fernández et al. [56] for *Thamnophis melanogaster*, Fathinia et al. [57] for *Pseudocerastes urarachnoides* and Sousa-Guedes et al. [58] for 13 different reptile species.

Apart from area expansion, our model also predicted an increase in the number of climatically suitable habitats in countries in which pancake tortoises do not exist naturally from the current two to eight future countries, with Angola isolated in the far west region (Fig 2; Table 1). The isolation of pancake tortoise populations is also occurring within Tanzania [42] and Kenya [10], where the species exists naturally. With pancake tortoises being non-migrant [10–12], this could be the reason for their non-existence in the climatically suitable habitats present in the other countries within the region. Furthermore, Malonza [10] has suggested that the absence of pancake tortoises in potential habitats is mainly due to elevation, with species occurring from 500 - 1,800 m above sea level [12]. As pancake tortoises prefer areas featuring Precambrian rocks, the presence of other rock types between the suitable habitats could act as a distribution barrier [13]. Therefore, this could be another reason for the non-existence of pancake tortoises in some climatically suitable habitats; however, the species may occur in the Zambezian floral region, provided that suitable habitat is available [13,37].

Surprisingly, pancake tortoises have been reported to occur naturally in Zambia [13] (Fig 1); however, our model did not indicate the presence of suitable habitats in Zambia either under current or future climatic scenarios (Fig 2; Table 1). One reason for this could be that the country is located at the limit of the climatically suitable niche and thus has low climatic suitability, which, when applied at a threshold, turns into an absence. This was affirmed by Chansa and Wagner [13], who have mentioned that the sightings of pancake tortoises in Zambia were recorded at the end of the pre-Cambrian rock basement. The absence of climatically suitable habitats for pancake tortoises in Zambia in current and future scenarios could mean that the animals recorded in the country were the result of the international animal trade, as a result of which animals from East Africa were exported illegally from the country [12]. However, this argument would require a genetic analysis for confirmation.

Our model also predicted that Tanzania, Kenya, Ethiopia and Angola (Fig 3; Table 1) will continue to have climatically stable habitats over time. As pancake tortoises have not yet been recorded in Ethiopia and Angola, these areas could hold potential for the translocation and introduction of the species. Additionally, we recommend studies of these countries to assess the existence of pancake tortoises in them, because the species is believed to exist in suitable habitats present in the Zambezian and Somalian biogeographical regions [37],[13].

Protected areas are one of the most necessary tools in biodiversity conservation [23]. However, the African Protected Areas Network does offer inconsistent protection to tortoise species [33]. In the Zambezian and Somalian biogeographical regions, only 32.37% of the current climatically suitable area for pancake tortoises fall within protected areas, and this percentage is predicted to decline in the future to 30.41% - 31.50% (Table 1). Additionally, from 66.92% - 85.13% of the stable climatic habitat is predicted to be outside of protected areas. Our results are inconsistent with those of Bombi et al. [33], who have suggested that the established protected areas in East Africa for wildlife conservation offer sufficient presentation for tortoises. Across the entire range, only 22.60% of pancake tortoise habitats are protected [33]. In Kenya, only 5.00% of the pancake tortoise population is protected, while in Zambia, none of the recorded species is within the Protected Area Network [12,13]. In Tanzania, only 4 out of 22 national parks have been recorded as harbouring pancake tortoises. That the pancake tortoise’s suitable habitat is largely unprotected in both the current scenario and the future increases the risk of overexploitation and habitat destruction as it has already been recorded in Tanzania [18], Kenya [10,12] and Zambia [12,13]. Additionally, the prevalence of ectoparasites such as ticks in the pancake tortoise is higher outside of the protected area [59]; this adds more risk of tick-borne diseases to the species.

### Management Implications

With the current and future climatically suitable habitats being found beyond the natural range of pancake tortoises, we recommend that studies be conducted in areas where pancake tortoises do not exist to confirm the occurrence of the species. As White [37] and Chansa and Wagner [13] have pointed out, pancake tortoises could exist in the entire Zambezian and Somalian biogeographical regions, provided that suitable habitat is present; therefore, confirmatory studies on the existence of the species in the climatically suitable habitats are essential for conservation planning for the species. However, we caution that the existence of pancake tortoises is not solely dependent on the presence of climatically suitable habitats, as Malonza [10] has confirmed the non-presence of pancake tortoises in typical habitats for the species in Kenya. Furthermore, the available suitable and stable habitats outside of the current range could be used as baseline areas for the translocation and introduction of the species where necessary. Therefore, we support the IUCN [19] and Bellis et al. [20], who have suggested the importance of conducting SDMs before translocation and species introduction/re-introduction. Our model did not predict the existence of climatically suitable habitats for pancake tortoises in Zambia (Fig 2). Therefore, we recommend the maximization of conservation efforts in Zambia in order to maintain the recorded pancake tortoise populations, since they seem to be highly threatened.

Furthermore, the presence of a large proportion of the climatically suitable habitat for pancake tortoises outside of protected areas could imply the need for more conservation efforts outside the protected range. These efforts might include the establishment of new protected areas aimed at biodiversity conservation to include suitable habitats for pancake tortoises and therefore minimize anthropogenic impacts on the species [10,35]. Since current increases to the Protected Area Network have rarely strategically considered global biodiversity maximization [23], establishing protected areas within species suitable habitats could be one strategy for protecting global biodiversity.

### Conclusion and Study Limitations

The SDM in this study predicted the expansion of suitable habitats for pancake tortoises in the future, which could lead to a stabilization of decreasing populational trends or even their inversion into a trend of growth. However, the largest proportion of these habitats will remain outside of the current Protected Area Network. This poses more risk to the species, considering it is critically endangered. Therefore, we do support the petition to upgrade the species from the current CITES Appendix II to Appendix I.

The findings of this study should not be treated as ready-made for on-the-ground application but could be used as one of many tools to help in conservation planning of the species, considering climatic changes. This is because our results were mainly based on climatic variables, and therefore, we did not consider non-climatic variables. This decision was influenced by the fact that most of the species’ distributional changes are largely driven by climatic variables [60]. Although Giannini et al. [61], de Araújo [6] and Palacio and Girini [61] have pointed out that the inclusion of biotic factors significantly improves SDMs, we were unable to obtain these data for our study. We recommend that future studies consider the inclusion of pre-Cambrian rock (as it provides a preferred habitat for pancake tortoises), the international pet trade, land-use changes and ecological interactions as variables. However, in the current situation, it is difficult to obtain these data ready-made for SDM, especially for future climatic scenarios. All in all, our study has provided the foundation for future studies on pancake tortoise distribution.

## ACKNOWLEDGEMENTS

We thank Tanzania Wildlife Research Institute (TAWIRI) and Commission for Science and Technology (COSTECH) for granting permit and Tanzania National Park (TANAPA) for giving free access into Tarangire National Park. We also thank Deo Tarimo for his help in obtaining pancake GPS locations from Mkomazi National Park and some from Tarangire National Park. Furthermore, we appreciate all the logistical support which were provided by the College of African Wildlife Management, Mweka during fieldwork.

## REFERENCES

1. Báez JC, Estrada A, Torreblanca D, Real R. Predicting the distribution of cryptic species: The case of the spur-thighed tortoise in Andalusia (southern Iberian Peninsula). Biodivers Conserv. 2012;21: 65–78. doi:10.1007/s10531-011-0164-3

2. Real R, Romero D, Olivero J, Estrada A, Márquez AL. Estimating How Inflated or Obscured Effects of Climate Affect Forecasted Species Distribution. PLoS One. 2013;8. doi:10.1371/journal.pone.0053646

3. Martínez-Freiría F, Argaz H, Fahd S, Brito JC. Climate change is predicted to negatively influence Moroccan endemic reptile richness. Implications for conservation in protected areas. Naturwissenschaften. 2013;100: 877–889. doi:10.1007/s00114-013-1088-4

4. Burrows MT, Schoeman DS, Richardson AJ, Molinos JG, Hoffmann A, Buckley LB, et al. Geographical limits to species-range shifts are suggested by climate velocity. Nature. 2014;507: 492–495. doi:10.1038/nature12976

5. Ceballos G, Ehrlich PR, Barnosky AD, García A, Pringle RM, Palmer TM. Accelerated modern human-induced species losses: Entering the sixth mass extinction. Sci Adv. 2015;1: 9–13. doi:10.1126/sciadv.1400253

6. Garcia RA, Cabeza M, Rahbek C, Araújo MB. Multiple dimensions of climate change and their implications for biodiversity. Science (80-). 2014;344: 1–10. doi:10.1126/science.1247579

7. Meng H, Carr J, Beraducci J, Bowles P, Branch WR, Capitani C, et al. Tanzania’s reptile biodiversity: Distribution, threats and climate change vulnerability. Biological Conservation. 2016. doi:10.1016/j.biocon.2016.04.008

8. Araújo MB, Alagador D, Cabeza M, Nogués-Bravo D, Thuiller W. Climate change threatens European conservation areas. Ecol Lett. 2011;14: 484–492. doi:10.1111/j.1461-0248.2011.01610.x

9. Raphael B, Klemens M, Moehlman P, Dierenfeld E, Karesh W. Blood values in free-ranging pancake tortoises (*Malacochersus tornieri*). J Zoo Wildl Med. 1995;25: 63–67.

10. Malonza PK. Ecology and Distribution of the Pancake Tortoise, *Malacochersus tornieri* in Kenya. J East African Nat Hist. 2003;92: 81–96. doi:10.2982/0012-8317(2003)92[81:eadotp]2.0.co;2

11. Mwaya RT, Moll D, Malonza PK, Ngwaya JM. *Malacochersus tornieri* (Siebenrock 1903) – Pancake Tortoise, Tornier’s Tortoise, Soft-shelled Tortoise, Crevice Tortoise, Kobe Ya Mawe, Kobe Kama Chapati. Chelonian Res Monogr. 2018;5: 107.1–107.15. doi:10.3854/crm.5.107.tornieri.v1.2018

12. Mwaya RT, Malonza PK, Ngwava JM, Moll D, Schmidt FA, Rhodin AGJN. *Malacochersus tornieri*. The IUCN Red List of Threatened Species 2019. IUCN Red List Threat. 2019. doi:https://doi.org/Malacochersus tornieri

13. Chansa W, Wagner P. On the status of *Malacochersus tornieri* (Siebenrock, 1903) in Zambia. Salamandra Bonn. 2006;42: 187.

14. Linder HP, de Klerk HM, Born J, Burgess ND, Fjeldså J, Rahbek C. The partitioning of Africa: Statistically defined biogeographical regions in sub-Saharan Africa. J Biogeogr. 2012;39: 1189–1205. doi:10.1111/j.1365-2699.2012.02728.x

15. Thulin M. Aspects of disjunct distributions and endemism in the arid parts of the Horn of Africa, particularly Somalia. Proceedings of the 13th Plenary Meeting, AETFAT, Zomba, Malawi. 1994. pp. 1105–1119.

16. Burgess N, Hales JD, Underwood E, Dinerstein E, Olson D, Itoua I, et al. Terrestrial Ecoregions of Africa and Madagascar. A Conservation Assessment. Island Press; 2004.

17. Moll D, Klemens MW. Ecological characteristics of the pancake tortoise, *Malacochersus tornieri*, in Tanzania. Chelonian Conserv Biol. 1996;2: 26–35. Available: https://scholar.google.com/scholar?hl=en&as_sdt=0%2C5&q=Moll+and+Klemens+%281996%29&btnG=

18. Klemens MW, Moll D. An assessment of the effects of commercial exploitation on the pancake tortoise, Malacochersus tornieri. Tanzania. Chelonian Conserv Biol. 1995;1: 197–206.

19. IUCN. Guidelines for Reintroductions and Other Conservation Translocations. Version 1.0. Gland, Switzerland: IUCN; 2013.

20. Bellis J, Bourke D, Maschinski J, Heineman K, Dalrymple S. Climate suitability as a predictor of conservation translocation failure. Conserv Biol. 2020. doi:10.1111/cobi.13518

21. White TH, de Melo Barros Y, Develey PF, Llerandi-Román IC, Monsegur-Rivera OA, Trujillo-Pinto AM. Improving reintroduction planning and implementation through quantitative SWOT analysis. J Nat Conserv. 2015;28: 149–159. doi:10.1016/j.jnc.2015.10.002

22. Craigie ID, Baillie JEM, Balmford A, Carbone C, Collen B, Green RE, et al. Large mammal population declines in Africa’s protected areas. Biol Conserv. 2010;143: 2221–2228. doi:10.1016/j.biocon.2010.06.007

23. Rodrigues ASL, Akçakaya HR, Andelman SJ, Bakarr MI, Boitani L, Brooks TM, et al. Global Gap Analysis: Priority Regions for Expanding the Global Protected-Area Network. Bioscience. 2004;54: 1092–1100. doi:10.1641/0006-3568(2004)054[1092:ggaprf]2.0.co;2

24. Coetzee BWT, Robertson MP, Erasmus BFN, van Rensburg BJ, Thuiller W. Ensemble models predict important bird areas in southern Africa will become less effective for conserving endemic birds under climate change. Glob Ecol Biogeogr. 2009;18: 701–710. doi:10.1111/j.1466-8238.2009.00485.x

25. Esser HJ, Mögling R, Cleton NB, Van Der Jeugd H, Sprong H, Stroo A, et al. Risk factors associated with sustained circulation of six zoonotic arboviruses: A systematic review for selection of surveillance sites in non-endemic areas. Parasites and Vectors. 2019;12: 1–17. doi:10.1186/s13071-019-3515-7

26. Warren R, VanDerWal J, Price J, Welbergen JA, Atkinson I, Ramirez-Villegas J, et al. Quantifying the benefit of early climate change mitigation in avoiding biodiversity loss. Nat Clim Chang. 2013;3: 678–682. doi:https://doi.org/10.1038/nclimate1887

27. Beaumont LJ, Graham E, Duursma DE, Wilson PD, Cabrelli A, Baumgartner JB, et al. Which species distribution models are more (or less) likely to project broad-scale, climate-induced shifts in species ranges? Ecol Modell. 2016;342: 135–146. doi:10.1016/j.ecolmodel.2016.10.004

28. Krechemer F da S, Marchioro CA. Past, present, and future distributions of bumble bees in South America: identifying priority species and areas for conservation. Journal of Applied Ecology. 2020. doi:10.1111/1365-2664.13650

29. Araújo MB, Williams PH, Fuller RJ. Dynamics of extinction and the selection of nature reserves. Hungarian Q. 2008;49: 1971–1980. doi:10.1098/rspb.2002.2121

30. Terribile LC, Lima-Ribeiro MS, Araújo MB, Bizão N, Collevatti RG, Dobrovolski R, et al. Areas of climate stability of species ranges in the Brazilian cerrado: Disentangling uncertainties through time. Nat a Conserv. 2012;10: 152–159. doi:10.4322/natcon.2012.025

31. Khan S, Nathand A, Das A. The distribution of the elongated tortoise (Indotestudo elongata)on the indian subcontinent: Implications for conservation and management. Herpetol Conserv Biol. 2020;15: 212–227.

32. Muñoz AR, Márquez AL, Real R. Updating Known Distribution Models for Forecasting Climate Change Impact on Endangered Species. PLoS One. 2013;8: 1–9. doi:10.1371/journal.pone.0065462

33. Bombi P, D’Amen M, Luiselli L. From Continental Priorities to Local Conservation: A Multi-Level Analysis for African Tortoises. PLoS One. 2013;8: 1–9. doi:10.1371/journal.pone.0077093

34. Villero D, Pla M, Camps D, Ruiz-Olmo J, Brotons L. Integrating species distribution modelling into decision-making to inform conservation actions. Biodivers Conserv. 2017;26: 251–271. doi:10.1007/s10531-016-1243-2

35. Church RL, Stoms DM, Davis FW. Reserve selection as a maximal covering location problem. Biol Conserv. 1996;76: 105–112.

36. UNEP-WCMC, IUCN. Protected Planet: The World Database on Protected Areas (WDPA), Cambridge, UK: UNEP-WCMC and IUCN. In: 2020 [Internet]. 2020. Available: www.protectedplanet.net

37. White F. The vegetation of Africa. A descriptive memoir to accompany the UNESCO-AETFAT-UNSO vegetation map of Africa, UNESCO, Paris. 1983; 356.

38. Coe MJ, Skinner JD. Connections, disjunctions and endemism in the eastern and southern african mammal faunas. Trans R Soc South Africa. 1993;48: 233–255. doi:10.1080/00359199309520273

39. Linder HP, Lovett J, Mutke JM, Barthlott W, Jürgens N, Rebelo T, et al. A numerical re-evaluation of the sub-Saharan phytochoria of mainland Africa. Biol Skr. 2005;55: 229–252.

40. Kabigumila J. Morphometrics of the Pancake tortoise (*Malacochersus tornieri*) in Tanzania. Tanzania J Sci. 2002;28: 34–46.

41. Wood RC, MacKay A. The distribution and status of the pancake tortoises, *Malacochersus tornieri*, Kenya. Conservafion, Restorafion and Management of Tortoises and Turtles-An Internafional Conference. 1997.

42. Spawls S, Howell K, Drewes R, Ashe J. A Field Guide to the Reptiles of East Africa - Kenya, Tanzania, Uganda, Rwanda and Burundi. San Diego, San Francisco, New York, Boston, London: AP Natural World; 2002.

43. Mwaya RT. The floristic composition of the habitat of *Malacochersus tornieri* at a hill in Tarangire National Park, Tanzania. Salamandra. 2009;45: 115–118.

44. Kyalo NS. Conservation, Management and Control of Trade in Pancake *Tortoise Malacochersus tornieri* (Siebenrock, 1903) in Kenya: The Non-Detriment Finding Studies Case Study. International Expert Workshop on CITES Non-Detriment Findings. Mexico,; 2008.

45. Chamberlain S, Barve V, Mcglinn D, Oldoni D, Desmet P, Geffert L, et al. rgbif: Interface to the Global Biodiversity Information Facility API. 2020. Available: https://cran.r-project.org/package=rgbif

46. Chamberlain S. rvertnet: Search “Vertnet”, a “Database” of Vertebrate Specimen Records. 2018. Available: https://cran.r-project.org/package=rvertnet

47. Iverson JB, Kiester AR, Hughes LE, Kimerling AJ. The EMYSystem world turtle database. In: http://emys.geo.orst.edu [Internet]. 2003 [cited 27 Apr 2020]. Available: http://emys.geo.orst.edu

48. Zacarias DA. Habitat analysis and population structure of the pancake tortoise (Malacochersus tornieri) in the Tarangire Ecosystem: an approach using GIS (Unpublished). College of African Wildlife Management, Mweka. 2007. Available: https://www.academia.edu/2563124/Aplicação_dos_SIG_e_análise_estatística_no_maneio_da_fauna_bravia_o_caso_das_tartarugas_de_carapaça_mole_Malacochersus_tornieri_

49. Karger DN, Conrad O, Böhner J, Kawohl T, Kreft H, Soria-Auza RW, et al. Climatologies at high resolution for the earth’s land surface areas. Sci Data. 2017;4: 1–20. doi:10.1038/sdata.2017.122

50. Sanderson BM, Knutti R, Caldwell P. A representative democracy to reduce interdependency in a multimodel ensemble. J Clim. 2015;28: 5171–5194. doi:10.1175/JCLI-D-14-00362.1

51. Naimi B, Hamm NAS, Groen TA, Skidmore AK, Toxopeus AG. Where is positional uncertainty a problem for species distribution modelling? Ecography (Cop). 2014;37: 191–203. doi:10.1111/j.1600-0587.2013.00205.x

52. R Core Team. R: A language and environment for statistical computing. Vienna, Austria. 2019. Available: https://www.r-project.org/

53. Naimi B, Araújo MB. Sdm: A reproducible and extensible R platform for species distribution modelling. Ecography (Cop). 2016;39: 368–375. doi:10.1111/ecog.01881

54. Barbet-Massin M, Jiguet F, Albert CH, Thuiller W. Selecting pseudo-absences for species distribution models: How, where and how many? Methods Ecol Evol. 2012;3: 327–338. doi:10.1111/j.2041-210X.2011.00172.x

55. Houniet DT, Thuiller W, Tolley KA. Potential effects of predicted climate change on the endemic south african dwarf chameleons, bradypodion. African J Herpetol. 2009;58: 28–35. doi:10.1080/21564574.2009.9635577

56. González-Fernández A, Manjarrez J, García-Vázquez U, D’Addario M, Sunny A. Present and future ecological niche modeling of garter snake species from the Trans-Mexican Volcanic Belt. PeerJ. 2018; 1–20. doi:10.7717/peerj.4618

57. Fathinia B, Rödder D, Rastegar-pouyani N, Hosseinzadeh MS, Kazemi SM. Zoology in the Middle East The past, current and future habitat range of the Spider-tailed Viper, *Pseudocerastes urarachnoides* (Serpentes: Viperidae) in western Iran and eastern Iraq as revealed by habitat modelling. Zool Middle East. 2020; 1–9. doi:10.1080/09397140.2020.1757910

58. Sousa-Guedes D, Arenas-Castro S, Sillero N. Ecological niche models reveal climate change effect on biogeographical regions: The Iberian Peninsula as a case study. Climate. 2020;8: 1–18. doi:10.3390/cli8030042

59. Mwaya RT, Mremi R, Eustace A, Ndibalema V. Prevalence of ticks (Acari: Ixodidae) parasitism on Pancake Tortoises, Malacochersus tornieri (Testudinidae), is lower inside than outside Tarangire National Park, Tanzania (In press). Chelonia Conserv Biol. 2020; Forthcoming.

60. Salas EAL, Seamster VA, Harings NM, Boykin KG, Alvarez G, Dixon KW. Projected future bioclimate-envelope suitability for reptile and amphibian species of concern in South Central USA. Herpetol Conserv Biol. 2017;12: 522–547.

61. Palacio FX, Girini JM. Biotic interactions in species distribution models enhance model performance and shed light on natural history of rare birds: a case study using the straight-billed reedhaunter Limnoctites rectirostris. J Avian Biol. 2018;49: 1–11. doi:10.1111/jav.01743

